# High-frequency oscillations reveal progressive recruitment of remote cortex into the epileptic network in a mouse model of focal cortical dysplasia type II

**DOI:** 10.64898/2026.09.03.748770

**Authors:** Nedime Karakullukcu, Michal Scheibel, Natalie Prochazkova, Bohdana Hruskova, Michaela Kralikova, Ondrej Novak, Helena Pivonkova, Jan Kudlacek, Premysl Jiruska

**Affiliations:** Department of Physiology, Second Faculty of Medicine, Charles University, Prague, 150 06, Czech Republic

**Keywords:** high-frequency oscillations (HFO), focal cortical dysplasia (FCD), epilepsy, interictal epileptiform discharges (IED), fast ripple, epileptogenesis

## Abstract

Interictal epileptiform discharges (IEDs) and pathological high-frequency oscillations - (HFOs) are established biomarkers of epileptogenic tissue, but how their spatiotemporal evolution reflects epileptic network reorganization in neocortical epilepsy remains unclear. We investigated longitudinal IED and HFO dynamics throughout epileptogenesis and chronic epilepsy in a mouse model of focal cortical dysplasia type II (FCD II). Long-term bilateral cortical recordings were obtained before and after spontaneous seizure onset. IEDs and HFOs (gamma, ripples, and fast ripples) were quantified in the dysplastic lesion and contralateral cortex during pre-epileptic, early, and late epileptic stages. The dysplastic lesion remained the principal seizure onset zone throughout disease progression and showed stable HFO activity after epilepsy onset. In contrast, the contralateral cortex exhibited progressive increases in IEDs and HFOs, demonstrating continuous recruitment into the epileptic network. Fast ripples were the earliest marker of this process, emerging within the first week after the first seizure and preceding increases in ripples, gamma, and IEDs. Propagation analysis showed that this contralateral increase was driven by independently generated HFOs rather than propagation from the lesion, indicating emergence of autonomous epileptogenic activity outside the primary focus. Spatiotemporal HFO analysis thus captures dynamic epileptic network remodeling beyond epileptogenic lesion. Fast ripples may serve as an early signature of network expansion and epileptogenicity emerging outside the primary lesion. Widespread structural and connectivity abnormalities extending beyond the lesion, combined with intense recurrent epileptic activity, may underlie the high endogenous epileptogenicity of FCD II, enabling small dysplastic lesions to recruit extensive neuronal networks across both hemispheres.

## 1. Introduction

High-frequency oscillations (HFOs) are brief (typically tens of milliseconds) epochs of oscillatory activity occurring in frequency bands above the conventional EEG range, generally spanning 80–800 Hz (Buzsáki et al., 1992; Worrell et al., 2004; Zijlmans et al., 2017). HFOs can be detected in both physiological and pathological conditions and are thought to reflect synchronized activity within local neuronal populations. HFOs span a wide frequency range, including ripple-band activity (80–250 Hz), which is important for normal brain function, including memory consolidation, sensorimotor processing, and movement execution (Bruder et al., 2021; Frauscher et al., 2018). Oscillations faster than ripples (>250 Hz) are considered mainly but not exclusively pathological HFOs (Frauscher et al., 2018). They typically emerge in hyperexcitable networks within aberrantly connected neuronal clusters, often co-occurring with EEG spikes (Liu et al., 2022; Ibarz et al., 2010; Frauscher et al., 2017). Such pathological HFOs, particularly fast ripples (FR, 250-500 Hz) have been reported in neurological disorders such as epilepsy or Alzheimer’s disease, highlighting their broader relevance in brain pathology (Worrell et al., 2004; Lisgaras et al., 2025). HFOs are used to localize epileptogenic tissue, the seizure onset zone, predict the future development of epilepsy, or monitor disease activity (Engel et al., 2013; Teurlings et al., 2026; Zijlmans et al., 2009; Jacobs et al., 2010). HFOs are strongly associated with regions of seizure initiation in the hippocampus and entorhinal cortex of patients with temporal lobe epilepsy (TLE), supporting the view that they reflect pathological hypersynchronous events crucially associated with epileptogenic tissue reorganization and its predisposition to generate spontaneous seizures (Staba et al., 2002). Much of the mechanistic understanding of pathological HFOs derives from TLE (Jacobs et al., 2009; Frauscher et al., 2017), where experimental and clinical findings have been mutually reinforcing (Jefferys et al., 2012). In neocortical epilepsy, clinical data on HFOs (Kerber et al., 2013) are not paralleled or extended by experimental research, largely because clinically relevant animal models have only recently become available (Lim et al., 2015). In particular, longitudinal studies that specifically link seizure evolution and interictal dynamics in neocortical epilepsy are missing.

In this study, we investigate how different HFO subtypes evolve during epileptogenesis and subsequent epilepsy progression in focal cortical dysplasia type II (FCD II). We show that HFOs, particularly fast ripples, provide information beyond that conveyed by IEDs about the progressive reorganization of FCD-related epileptic networks. We demonstrate that epileptic networks in FCD II progressively expand beyond the primary lesion, a process that may be facilitated by pre-existing morphological, connectivity, synaptic, and intrinsic abnormalities associated with FCD II. These abnormalities may create a structurally and functionally permissive substrate ultimately promoting the emergence of epileptogenicity outside the primary lesion.

## 2. Methods

### 2.1. Animal Model

All experiments were performed under the Animal Care and Animal Protection Law of the Czech Republic, fully compatible with the guidelines of the European Union directive 2010/63/EU. The protocol was approved by the Ethics Committees of the Second Faculty of Medicine (Project License No. MSMT-31765/2019–4). Experimental data from 8 mice were analyzed in this study (5 females and 3 males). Three were wild-type C57BL/6N mice, and three were NG2-DsRed reporter mice (Tg(Cspg4-DsRed.T1)1Akik/J; JAX no. 008241, used as heterozygous = +/-) mice back-crossed to the C57BL/6N background. In addition, one animal was of B6.129-Gt(ROSA)26Sortm1(CAG-CHRM4*,-mCitrine)Ute/J (Jackson lab. no.026219) strain with cre-dependent expression of hM4Di receptor and one animal was generated by crossing SST-IRES-Cre (Cre recombinase expressed under the somatostatin promoter; Jackson Laboratory, no.018973) mice with Ai14 reporter mice (Cre-dependent expression of the red fluorescent protein tdTomato; Jackson Laboratory, no.007909). The mice were used also in other studies and therefore, the cohort has a heterogeneous background. Animals were housed under 12/12 light-dark cycles with ad libitum access to food and water, were bred in house, and had undergone no experimental procedure before the electrode implantation reported here: no cranial window implantation, no viral vector injection, no chemogenetic or optogenetic manipulation, no imaging procedure and no pharmacological treatment. The FCD induction, surgeries and EEG recording were performed according to Chvojka et al. (2024). Briefly, all eight mice underwent FCD induction by *in utero* electroporation of a plasmid containing mutated mTOR (p.Leu2427Pro; SoVarGen Co., Ltd., Korea) and fluorescent reporter protein (GFP or iRFP) to locate the position of FCD lesion. Another FCD animal was generated by *in utero* electroporation of cre-GFP-T2A-HA-RhebCA (in house designed, produced by VectorBuilder) and iRFP.

The study used a single-group, within-subject longitudinal design. All eight animals carried a dysplastic lesion and received the same intervention, and each animal was compared with its own pre-epileptic period. For this reason, no separate non-lesioned control group was recorded. The individual animal was the experimental unit in all inferential analyses.

### 2.2. Signal Acquisition

At the age of 8 weeks, mice were implanted 4 or 5 epidural electrodes positioned over the frontal and parietal cortices of the lesional hemisphere (lesional, L; peri-lesional, PL; frontal left, FL) and the corresponding contralateral hemisphere (frontal right, FR; contralateral, C) and a ground/reference electrode over the cerebellum. The EEG was recorded using Intan RHD system (Intan Technologies, USA) at a sample rate of 5 kHz continuously for >3 weeks. Seizures were identified visually by observers who were not involved in the design or the analysis of the present study and who were unaware of its hypotheses, and the day of the first spontaneous seizure was set as 0. The age at first spontaneous seizure was 64.9 ± 14.7 days (mean ± SD; range 45.2-86.3 days). Since the temporal distribution and the day of the first seizure differ between animals, we have decided to reference the longitudinal analyses to the first seizure rather than age of the mice or other artificial time frame.

### 2.3. Spike and HFO Detection

We implemented the IED and HFO detection framework in MATLAB. The pipeline is shown in Supplementary Figure 1. IEDs were detected using a Hilbert envelope-based approach (Janca et al., 2015). Unfiltered IEDs were used as the input to the HFO detection pipeline. HFOs were identified using a detector adapted from Cho et al. (2014). Signals were first processed using a band-pass filter followed by the Hilbert transform to extract the analytic amplitude and instantaneous frequency. Band-pass filtering was performed using a linear-phase finite impulse response (FIR) filter (order = 150) applied to data sampled at 5 kHz. Following the filtering, candidate events were identified using an amplitude and duration thresholding procedure (Cho et al., 2014). Rectified signals were analyzed to compute the cycle count; a minimum of 6 peaks was required for the detection of putative HFO events. The output of this stage was a set of initial HFO candidates.

Each putative HFO event underwent multi-domain feature extraction, including kurtosis, skewness, Shannon entropy, log energy entropy, energy, discrete wavelet transform (DWT) coefficients, power spectral density (PSD) estimate such as delta-band power (0.5–4 Hz), theta-band power (4–8 Hz), the delta/theta power ratio, alpha-band power (8–12 Hz), beta-band power (13–30 Hz), the alpha/beta power ratio, combined gamma+ripple power (30–250 Hz), fast-ripple power (250–800 Hz), the ratio of combined gamma+ripple to fast-ripple power, all-band power (0.5–1000 Hz), and autoregressive (AR) coefficients (order = 4). In total, 17 features were extracted. A brief description of each feature and its mathematical formulation is provided in Supplementary Table 1. Based on these features, each putative HFO event was classified as HFO or non-HFO using a trained ensemble bagging model (Supplementary Fig. 2). Classifier training, cross-validation, and performance are detailed in Supplementary Methods and Supplementary Results (Supplementary Fig. 3). Events classified as HFO were further categorized into three frequency bands: gamma (30–80 Hz), ripple (80–250 Hz) or fast ripple (250–800 Hz) median of inter-spike intervals within the event (*ISI_med_*) according to Table 1 (Jiruska et al., 2010).

**Table 1:**
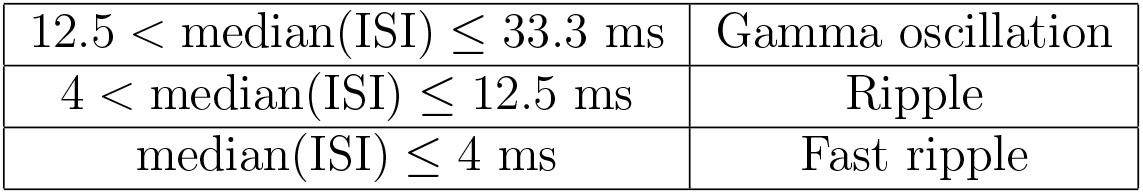
Classification of HFO events into frequency bands.

For each animal, the total recording duration was computed by summing all valid recording signal. Event rates were computed as the number of events divided by the total duration of valid signal, thus taking into account possible signal dropouts or artifact contamination. Therefore, periods were not biased by circadian sampling.

### 2.4. HFO Spectral Properties and Network Propagation

To characterize the temporal profile of interictal events, we analyzed three periods referenced to the first seizure: i) pre-epileptic period (day −7 to 0), ii) early-epileptic period (day 0 to 7), and iii) late-epileptic period (day 14 to 20). To characterize whether the progressive increase in HFO activity is accompanied by changes in signal spectral content and network organization, we analyzed power spectral density (PSD) of IED-coupled HFO subtypes and the spatial propagation patterns of individual subtypes across the three temporal periods. To characterize the broadband spectral properties of IEDs coupled with HFOs, we quantified the PSD of events detected by the automated algorithm. Because the dataset contained an exceptionally large number of events (>39,000 fast ripples), we employed a fixed-size random subsampling procedure to achieve computational efficiency while maintaining statistical representativeness. For each animal, period, and hemisphere, up to 1000 HFO-coupled IEDs were randomly selected using a fixed seed of random number generator to ensure full reproducibility. PSD estimation was performed on 300-ms segments centered on each selected event using Welch’s method (pwelch; 50% overlap, Hann window 128 ms). Before computing the spectra, all segments belonging to the same period and hemisphere were grouped, and signals underwent a standardized preprocessing pipeline that included linear detrending, notch filtering at 50 and 100 Hz, artifact rejection based on a median absolute deviation (MAD) threshold (5 × MAD), and Hann windowing. Segments within each group were then concatenated, and a single power spectral density estimate was computed using Welch’s method (pwelch). PSD estimates were finally averaged across animals, separately for lesional and contralateral channels.

To analyze HFO propagation between hemispheres and its long-term evolution, we categorized the events as follows. HFO onset was defined as the initial point of the binary detection signal (the first sample at which the rectified signal’s RMS envelope exceeded the mean envelope + 2.5 x SD, computed over a 2 ms sliding window). Inter-hemispheric propagation delays were then computed as the difference between the lesional (L) and contralateral (C) onset times for events co-occurring within a ±50 ms window. If an event in one hemisphere was followed by the same type of event in the other hemisphere with a delay 1.5 *<* Δ*t <* 50 ms, it was labeled as L→C or C→L, according to the sign of the delay. If Δ*t <* 1.5, the event was deemed to occur simultaneously with no propagation and was excluded from the propagation analysis. Events separated by more than 50 ms were regarded as individual events. The propagation rates were normalized by the rates of the given event type.

### 2.5. Statistical Analysis

Statistical analyses were performed using MATLAB (version R2024b, MathWorks, Natick, MA, USA). For each animal, we computed event rates in 1-hour time bins. Bins containing seizures were excluded. Given the limited sample size (*n* = 8), animals with different genetic backgrounds were analyzed together, and neither genotype nor sex was included as an independent factor in the statistical analyses. Animals constituted the independent experimental units in all inferential analyses. Measurements obtained across days and recording channels were treated as repeated observations nested within animals and were accounted for using paired non-parametric tests or linear mixed-effects models (LME) with animal identity specified as a random effect. Consequently, days relative to the first spontaneous seizure were not treated as independent observations, thereby avoiding pseudoreplication. Channel-specific analyses were used to describe the spatial localization of activity.

The Wilcoxon signed-rank comparisons of event rates between periods and between channels were applied separately for each event type. Similarly, the Wilcoxon signed-rank comparisons of band-limited power between periods and between channels were applied separately for each frequency band. To assess time-dependent changes, LME were fitted across all 1-hour bins, with event rate as the dependent variable, for each metric with day to seizure as a fixed effect and animals as a random effect. The estimated slope therefore, represents the average change in event rate per day (events/h/day). The significance of the temporal trend was assessed using the *p*-*value* of the fixed-effect coefficients. Additionally, for the day-by-day course of statistical significance over the 20 days following the first seizure, we averaged the rates from recording start to the first seizure onset, excluding the last 6 hours before the first seizure onset, and used them as a baseline. After that, we compared the baseline with each day after the first seizure using LME. Resulting p-values were corrected for multiple comparisons across the 20 tested days within each event-rate metric using the Benjamini-Hochberg false discovery rate (FDR) procedure, applied separately for each metric (IED, gamma, ripple, and fast ripple rate)

For the propagation analysis, the statistical significance of changes across pre-epileptic, early-epileptic, and late-epileptic periods was evaluated using the Friedman test.

Effect sizes were classified using Cohen’s (Cohen, 1988) thresholds. For Wilcoxon signed-rank tests the effect size r was computed as:

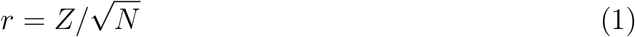

where r is the magnitude of the paired difference, Z is the test statistic from the normal approximation of the signed-rank test, and N is the number of paired observations (*trivial <* 0.10, *small* 0.10–0.30, *medium* 0.30–0.50, *large* ≥ 0.50). For the linear mixed-effects models, the fixed-effect slope with its 95% confidence interval is reported as the measure of effect size. We used Kendall’s coefficient of concordance (*W*) to compute effect size for the Friedman test with the formula:

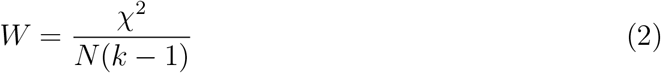

where *X*^2^ is the Friedman test statistic, *k* and *N* are the number of groups and the total number of animals, respectively. *p*-values were adjusted for multiple comparisons using the Benjamini–Hochberg false discovery rate (FDR) procedure to account for multiple comparisons across features and tests. Adjusted *p*-value < 0.05 were considered statistically significant. All results and graphs are shown as mean ± SD (median ± IQR) unless otherwise stated.

## 3. Results

We analysed data from eight mice that met the a priori criterion of a recording window spanning at least 2 days before the first recorded seizure. This seizure was then considered the first seizure the mouse experienced and the onset of epilepsy. Within these windows, the duration of valid, artefact-free signal after exclusion of dropouts ranged from 1.7 to 25.9 days (9.5 ± 8.5 (7.3 ± 12.2)). After the first seizure, we used at least 3 weeks of recording for 7 mice and 10 days for one animal. The data set contained 398 seizures. Across the eight FCDII mice, the timing and burden of spontaneous seizures varied considerably (Fig. 1). Five subjects (Mouse02, Mouse03, Mouse06, Mouse07, Mouse08) displayed prolonged seizure-free intervals followed by seizure clusters, whereas 3 animals (Mouse01, Mouse04, Mouse05) exhibited relatively static seizure rate as described previously (Kudlacek et al., 2025).

**Figure 1:**
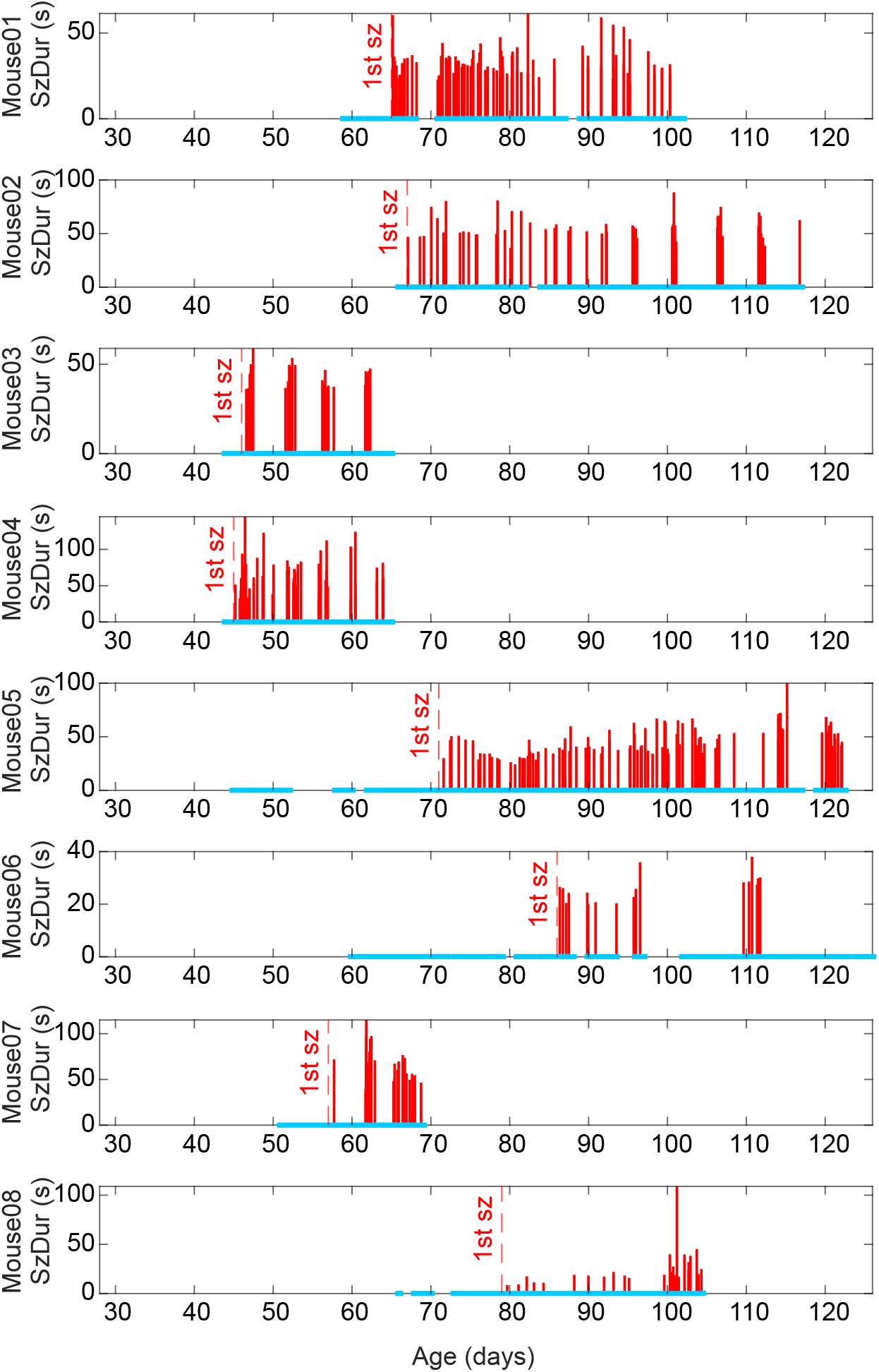
Seizure profiles across individual subjects. Raster plots show the temporal distribution and duration of seizures (SzDur; red bars) for each subject over the monitoring period. The blue baseline indicates the presence of recording, gaps represent recording dropouts. The dashed vertical line marks the time of the first recorded seizure (1st sz) for each animal.

During the observation period of 44.2 ± 23.2 (44.1 ± 39.9) days, each mouse had 49 ± 31 (44 ± 47) seizures, and the mean seizure frequency was 1.34 ± 0.82 (1.28 ± 1.14) seizures per day, and seizure duration was 45.2 ± 16.7 (44.2 ± 30.3) s. To determine the main seizure onset zone, for each seizure, cross-correlation between each pair of channels was computed within a ±1 s window around seizure onset. The channel exhibiting the most negative lag from any of the other channel at maximum correlation was defined as the earliest activated channel. Using this approach, we revealed that 95.7 ± 5.2 (96.6 ± 6.0)% of seizures originated from lesional or nearby channels (L, PL, FL; data not shown). We also compared seizure onsets during the early and late stages of epilepsy. The comparison of the seizure onset zone between the early- and late-epileptic periods was restricted to six animals. One animal had no recording during the late-epileptic period, because its recording ended 11 days after the first seizure, and one further animal had no seizures during the late-epileptic period, so no onset zone could be determined for that window. The results showed that the lesion hemisphere was the predominant seizure onset zone in both stages (early: 67.9 ± 32.4% (69.4 ± 56.2%); (late: 87.4 ± 19.5%, (95.5 ± 16.7%)), with no difference between stages (Wilcoxon signed-rank, *p* = 0.375, *n* = 6 subjects). The lesion-to-contralateral (L→C) propagation lag computed over the first 5 seconds of seizure was shifted from positive to negative from the stages early-epileptic to late-epileptic (early: +0.04 ± 0.65 ms; late: −0.08 ± 0.86 ms; *p* = 1.000), indicating that the contralateral cortex was activated after the first seizure onset, with no significant change across disease stages. Interictal periods were characterized by the presence of various forms of interictal epileptiform discharges (IEDs, single spikes, polyspikes) and pathological HFOs from gamma, ripple, and fast ripple bands (Fig. 2). In total, 2,213,905 IEDs were detected, corresponding to an average rate of 5.70 ± 2.57 (6.65 ± 3.68) IEDs/min. Superimposed on them, we detected 5,168 gamma events, occurring at rate of 0.015 ± 0.012 (0.014 ± 0.015) events/min, 3,414 ripples occurring at rate 0.010 ± 0.008 (0.008 ± 0.010) events/min, and 39,547 fast ripples occurring at rate of 0.11 ± 0.07 (0.10 ± 0.135) events/min.

**Figure 2:**
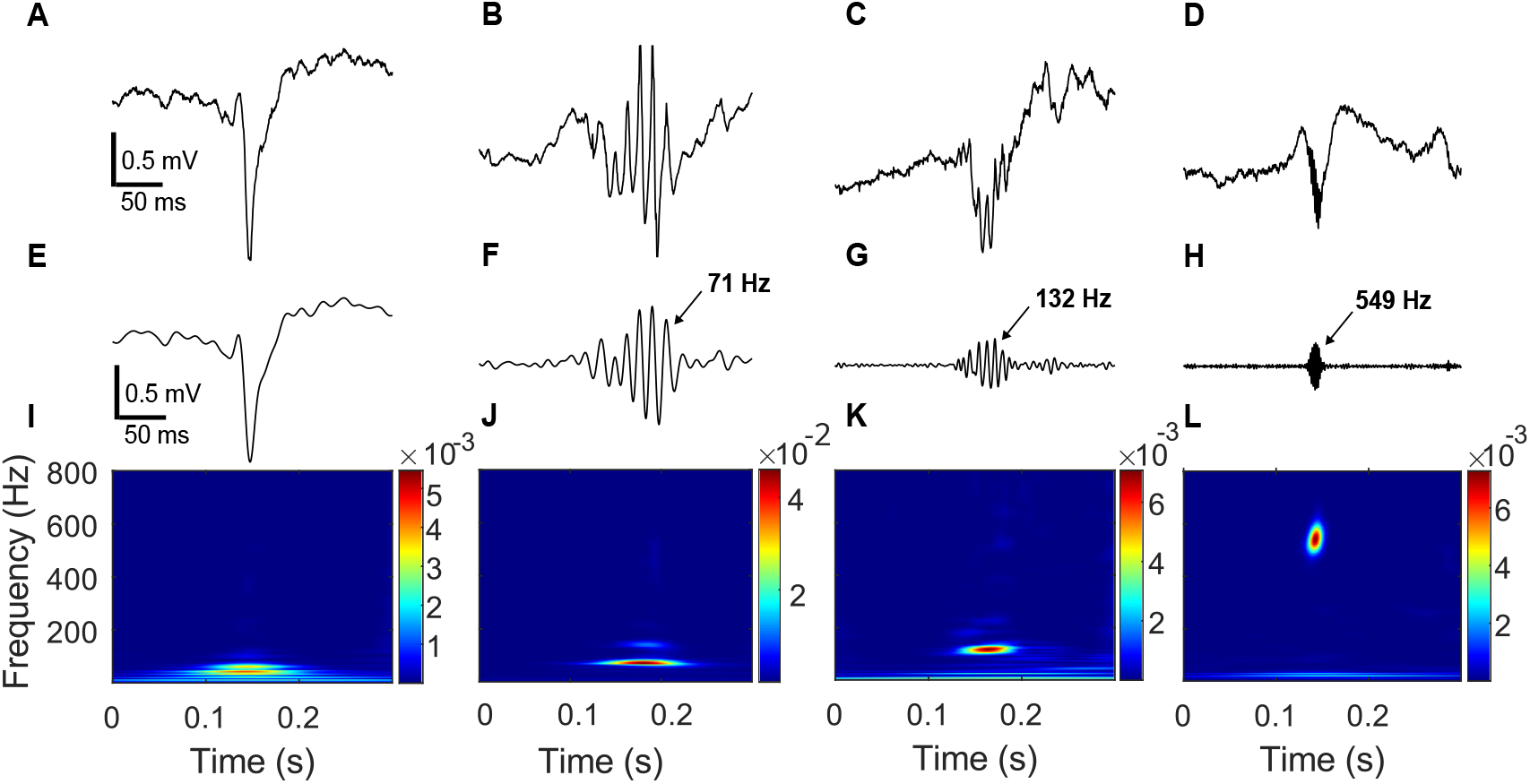
Example IED and HFO subtypes. Raw EEG signals (A–D) and corresponding bandpass-filtered signals (E–H) in the IED (0.5–80 Hz), gamma (30–80 Hz), ripple (80–250 Hz), and fast ripple (250–800 Hz) ranges are shown. Time-frequency representations (I–L) were computed for the full-band signal (0.5–800 Hz) using superlet transform (Moca et al., 2021). Amplitude and frequency scales were kept identical across events to facilitate direct comparison.

### 3.1. Channel Specific Analyses of IEDs and HFOs

In this study, we examined the spatiotemporal profiles of IEDs and HFOs recorded from electrodes in the FCD lesion (L) and the contralateral homotopic neocortex (C). The temporal and spectral evolution of IED and HFO subtype activities in the FCD model across three disease phases: pre-epileptic (−7 to 0 d), early-epileptic (0-7 d), and late-epileptic (14-20 d) (Fig. 3). We quantified the hourly burden of IEDs and HFO subtypes in the lesion vs the contralateral hemisphere (Fig. 3A-D) (Wilcoxon signed-rank tests, FDR-corrected across all pairwise comparisons; r = rank-biserial effect size). In the FCD lesion, only IEDs showed a change in the rate of occurrence (Fig. 3A); there was an increase from the pre- to early-epileptic period (*n* = 8), early- to late-epileptic (*n* = 7), and pre- to late-epileptic (*n* = 7)(in all cases *p* = 0.016, *r* = 0.94). Contralateral IED rates did not change significantly across any period transition (*p >* 0.14), yet they were consistently higher than lesional rates at every period (lesion vs. contralateral: pre-epileptic (*p* = 0.035, *r* = 0.89, *n* = 8), early-epileptic (*p* = 0.039, *r* = 0.83, *n* = 8), and late-epileptic (*p* = 0.035, *r* = 1.00, *n* = 7). The HFO subtypes behaved differently, with significant changes in HFO rates appearing only on the contralateral side. Gamma and ripple rates in the lesion were stationary, while their rates in the contralateral hemisphere increased from the pre-epileptic to late epileptic period (Fig. 3B and C; *p* = 0.045, *r* = 1.00 in both). In the contralateral hemisphere, fast ripple rate increased from the pre-epileptic period to both the early- (Fig. 3D; *p* = 0.035, *r* = 0.89, *n* = 8) and late-epileptic phases (*p* = 0.035, *r* = 1.00, *n* = 7). The lesional cortex showed no comparable change in fast ripple rates. Because HFOs are a more reliable biomarker of endogenous epileptogenicity than IEDs (Chvojka et al., 2024), these longitudinal HFO-rate profiles suggest that the contralateral cortex undergoes more profound reorganization than the lesion following the first seizure. To validate the observed changes in HFO rates, we also performed spectral analysis of detected IEDs during individual stages. In the lesion, no band showed a statistically significant band power change from the pre- to late-epileptic phase (Wilcoxon signed-rank, all *p >* 0.23), despite a non-significant increasing trend in fast ripple power in a subset of animals. Conversely, in the contralateral cortex, gamma and fast ripple band power significantly increased from the pre-epileptic to late-epileptic phase (*p* = 0.047, *r* = 1.00, *n* = 7 for both bands), supporting progressive contralateral hemisphere recruitment, which is not paralleled by similar reorganization in the lesion. Consistent with this, lesion and contralateral power differed significantly in the gamma and ripple bands specifically during the early-epileptic phase (Fig. 3F; both *p* = 0.023, *r* = 0.94, *n* = 8), but not during the pre- or late-epileptic phases (all *p >* 0.16), indicating a transient lesion–contralateral cortex asymmetry that resolves as the contralateral hemisphere becomes progressively recruited (Fig. 3E-G).

**Figure 3:**
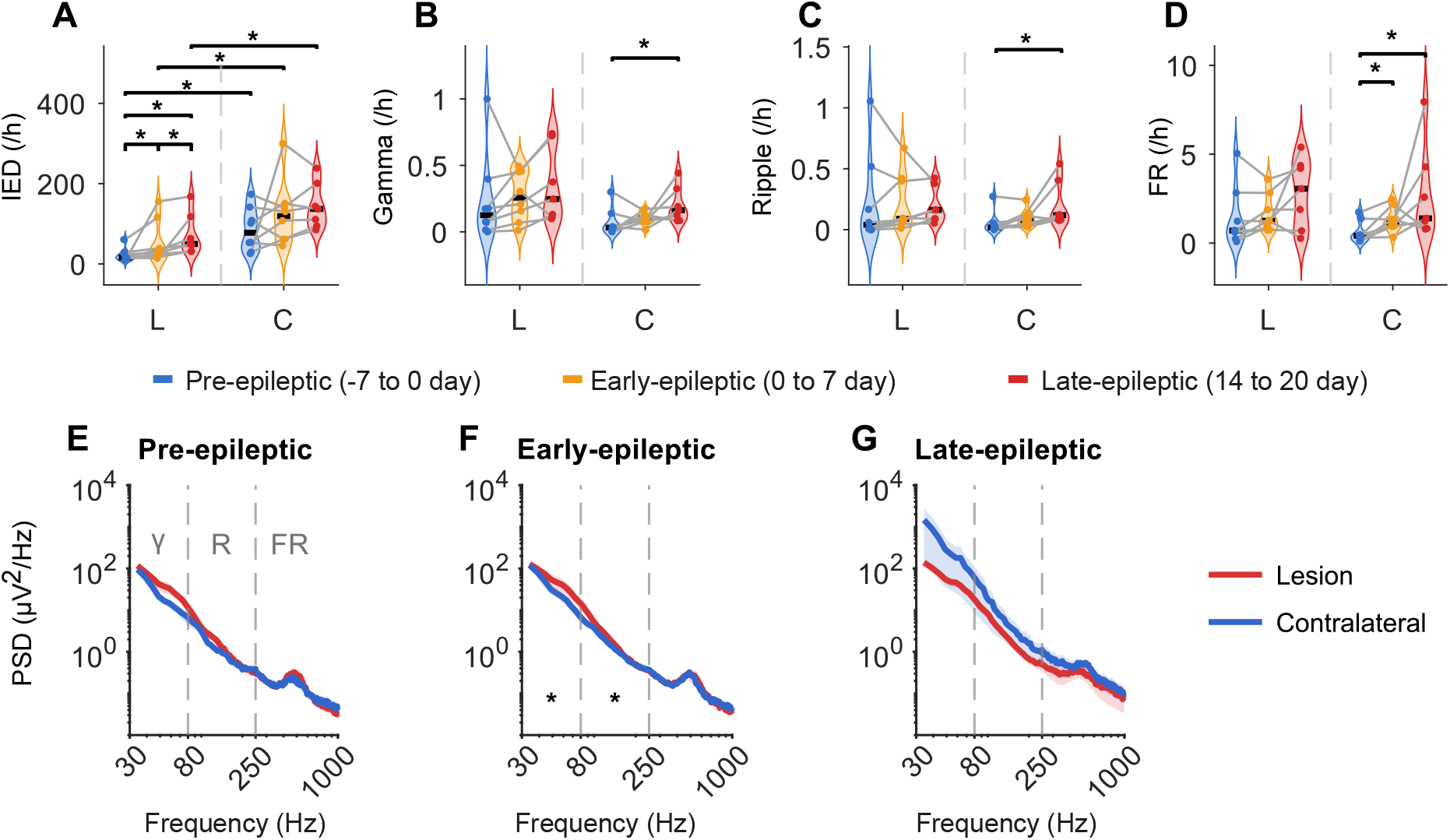
The temporal and spectral evolution of IEDs and HFO subtypes across three disease phases. (A–D) Event rates per hour for IED and HFO subtypes in lesional, L, and contralateral, C, channels. Data points from individual animals are connected. In the lesion, only the IED rate increased across every period transition, whereas none of the HFO subtypes changed. The contralateral channel showed the opposite pattern: the IED rate did not change across periods but exceeded the lesional rate at every period, while gamma and ripple rates increased from the pre- to the late-epileptic period and fast ripples (FR) increased from the pre-epileptic period to both subsequent periods. (E–G) PSD of IED-associated activity in lesion (red) and contralateral (blue) channels during the pre-, early-, and late-epileptic periods; thin lines show individual animals, thick lines show the group mean. (Wilcoxon signed-rank test * *p <* 0.05.)

Next, we performed a detailed longitudinal analysis of the IED and HFO rate evolution in each hemisphere (Figs. 4 and 5). One of the eight animals was excluded from these analyses because its recording terminated too early (11 days after the first seizure). We first plotted the long-term group-level traces showing the raw time course for each event from seven days before to twenty days after the first seizure in the lesional channel in Fig. 4A. To determine not only whether, but precisely when, event rates diverged from each animal’s own pre-ictal baseline, we additionally compared the rate on each of the 20 days following the first spontaneous seizure to that animal’s baseline using linear mixed-effects models. We mapped the day-by-day course of statistical significance over the 20 days following the first seizure using a thickened line with a highlighted transparent gray bar. Significance here was evaluated using a linear mixed-effects (LME) model with FDR correction. In the lesion, only the IED rate was different from the baseline on the 20th day only (*p* = 0.0017, *n* = 7). LME models applied to the whole recordings revealed significant upward temporal trends in the lesional channel for IED (Fig. 4B, slope = 3.11 (events/h, per day), 95% CI [0.99, 5.31], *p* = 0.0054, *n* = 7), gamma (Fig. 4C, slope = 0.01 (events/h, per day), 95% CI [0.0036, 0.011], *p* = 0.00013, *n* = 7), and fast-ripple rates (Fig. 4E, slope = 0.07 (events/h, per day), 95% CI [0.01, 0.14], *p* = 0.023, *n* = 7). Ripple rate (Fig. 4D, slope = 0.00 (events/h, per day), 95% CI [-0.01, 0.008], *p* = 0.81, *n* = 7) showed no significant trend.

**Figure 4:**
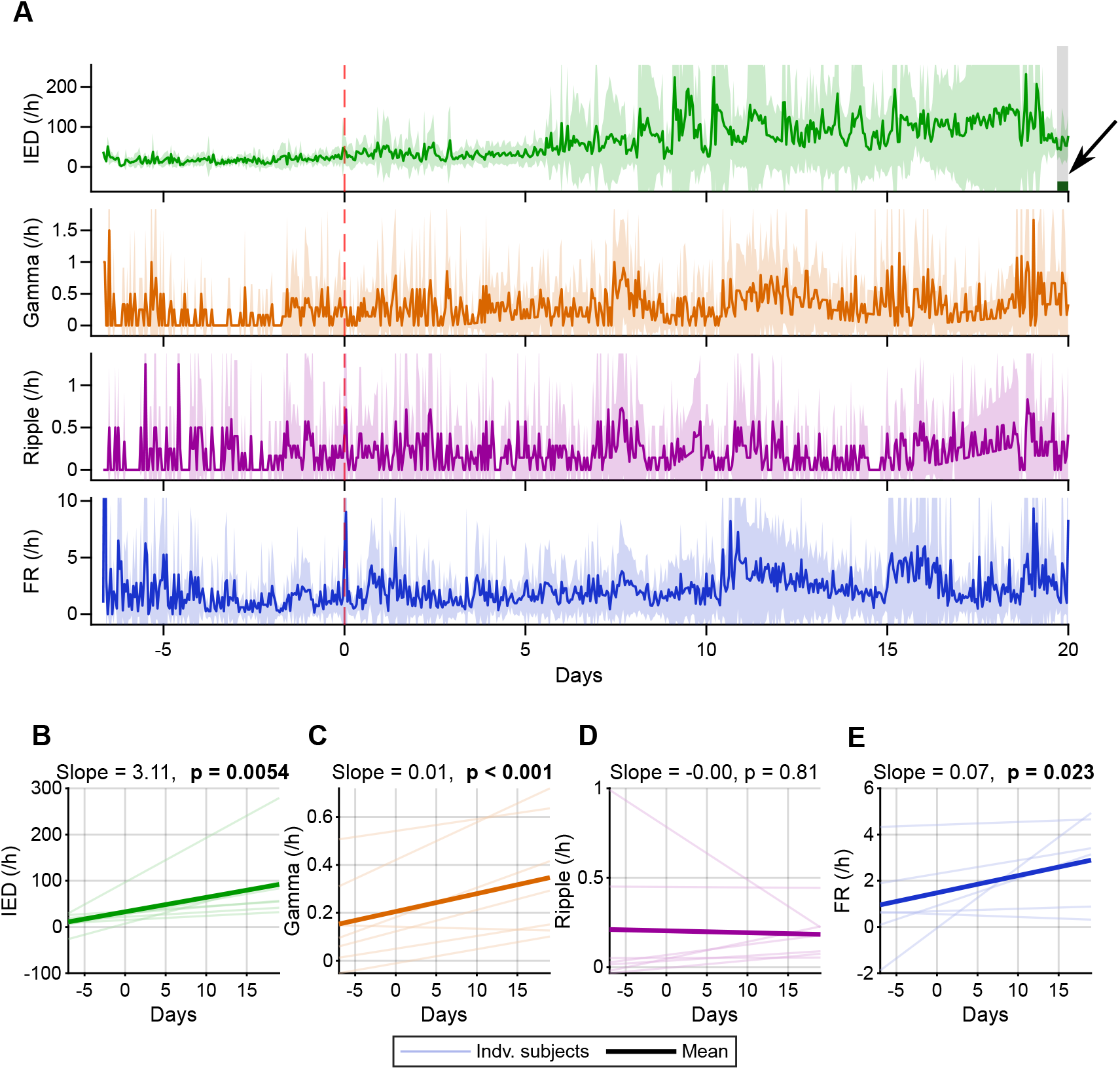
Temporal evolution of event rates in the lesional channel (n = 7 animals). (A) Group-mean hourly rates and (± SD, shaded) of IEDs, gamma, ripple, and fast ripple (FR), aligned to the first spontaneous seizure. Colored bars with a highlighted transparent gray rectangle on each trace mark the days on which the event rate differed significantly from baseline (LME, FDR-corrected). Only the IED rate differed from baseline, and on a single day (day 20), which does not demonstrate a discrete change in lesional activity at the onset of spontaneous seizures. (B-E) Linear mixed-effects model fits for IEDs and HFO subtype event rates against time. Slope values indicate the mean change in the units of event rate (e.g. IED/h) per day. The rates of IEDs (B), gamma (C) and fast ripples (E) increased significantly, whereas the ripple rate (D) did not change. Thin pale lines represent individual subjects; thick colored lines indicate group means.

**Figure 5:**
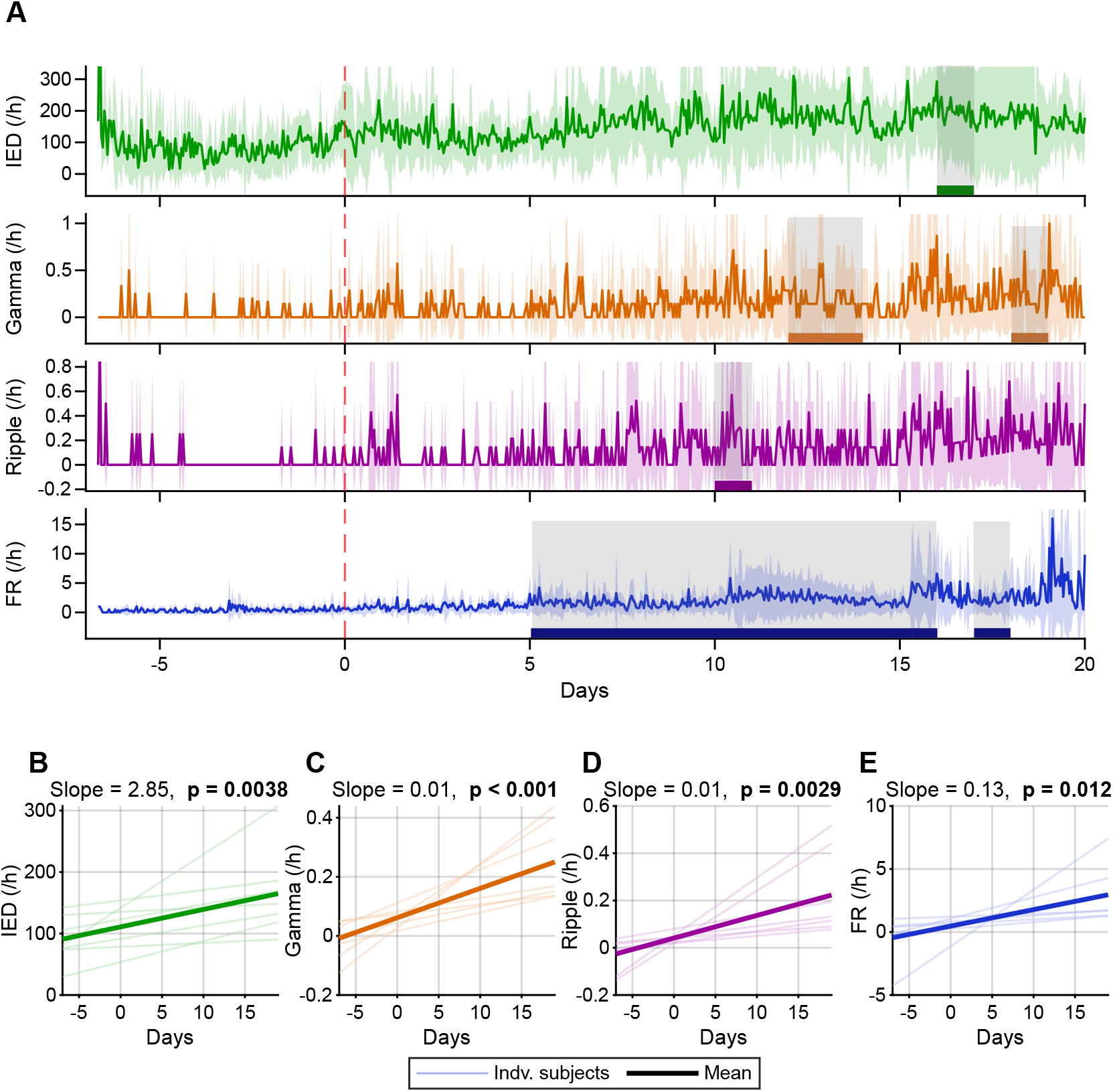
Temporal evolution of event rates in the contralateral channel. (A-E) Plotting conventions and statistics as in Fig. 4. (A) Fast ripples (FR) were the first subtype to diverge from baseline, followed by ripples on day 10, gamma on days 12, and IEDs on day 16. Colored bars with a highlighted transparent gray rectangle on each trace mark the days on which the event rate differed significantly from baseline (LME, FDR-corrected). The rates of IEDs (B), gamma (C), ripple (D) and fast ripples (FR; E) all increased significantly.

In contrast, the contralateral channel showed a significant increase in the IED or HFO rates compared to baseline on multiple days, which is indicated by the thick line at the bottom of each graph (Fig. 5A). IED rates reached significance only on day 16 (*p* = 0.014, *n* = 7). Gamma rates showed two discrete windows of significance, one around days 12-13 (*p* = 0.04, *p* = 0.0055, and *n* = 7) and a second day 18 (*p* = 0.04, *n* = 7). Ripple rates shifted later, crossing the significance threshold around day 10 (*p* = 0.032, *n* = 7). Fast ripple activity showed the earliest changes, reaching significance the 5th day after the first seizure (*p* = 0.013, *n* = 7), and staying significant through day 15, with a second window of significance re-emerging around days 17. All four metrics showed significant positive temporal trends by LME: IED (Fig. 5B, slope = 2.85 (events/h, per day), 95% CI [0.92, 4.77], *p* = 0.0038, *n* = 7), gamma (Fig. 5C, slope = 0.01 (events/h, per day), 95% CI [0.0051, 0.015], *p* = 5.4 *×* 10*^−^*^5^, *n* = 7), ripple (Fig. 5D, slope = 0.01 (events/h, per day), 95% CI [0.0033, 0.016], *p* = 0.0029, *n* = 7), and fast ripple (Fig. 5E, slope = 0.13 (events/h, per day), 95% CI [0.028, 0.23], *p* = 0.012, *n* = 7). Together, these results suggest that while the lesional cortex maintains a chronically elevated but relatively stable activity level, the contralateral cortex undergoes progressive and time-dependent reorganization following the first spontaneous seizure.

### 3.2. Inter-hemispheric Propagation and Network Reorganization of HFOs

After demonstrating progressive involvement of contralateral cortex in FCD-related epileptic network, we aimed to answer the question whether the involvement is attributed to the facilitated propagation of lesional discharges and HFOs or whether the contralateral cortex becomes an independent generator of IEDs and HFOs. To quantify the propagation of events between hemispheres, we computed the rate of fast ripple events propagating from the lesional to the contralateral hemisphere (L→C) and from the contralateral to the lesional hemisphere (C→L) for the three analysed periods. Examples of L→C, C→L propagating, isolated in the L-only and isolated in the C-only fast ripples are shown in Fig. 6. More examples are in Supplementary Fig. 4.

**Figure 6:**
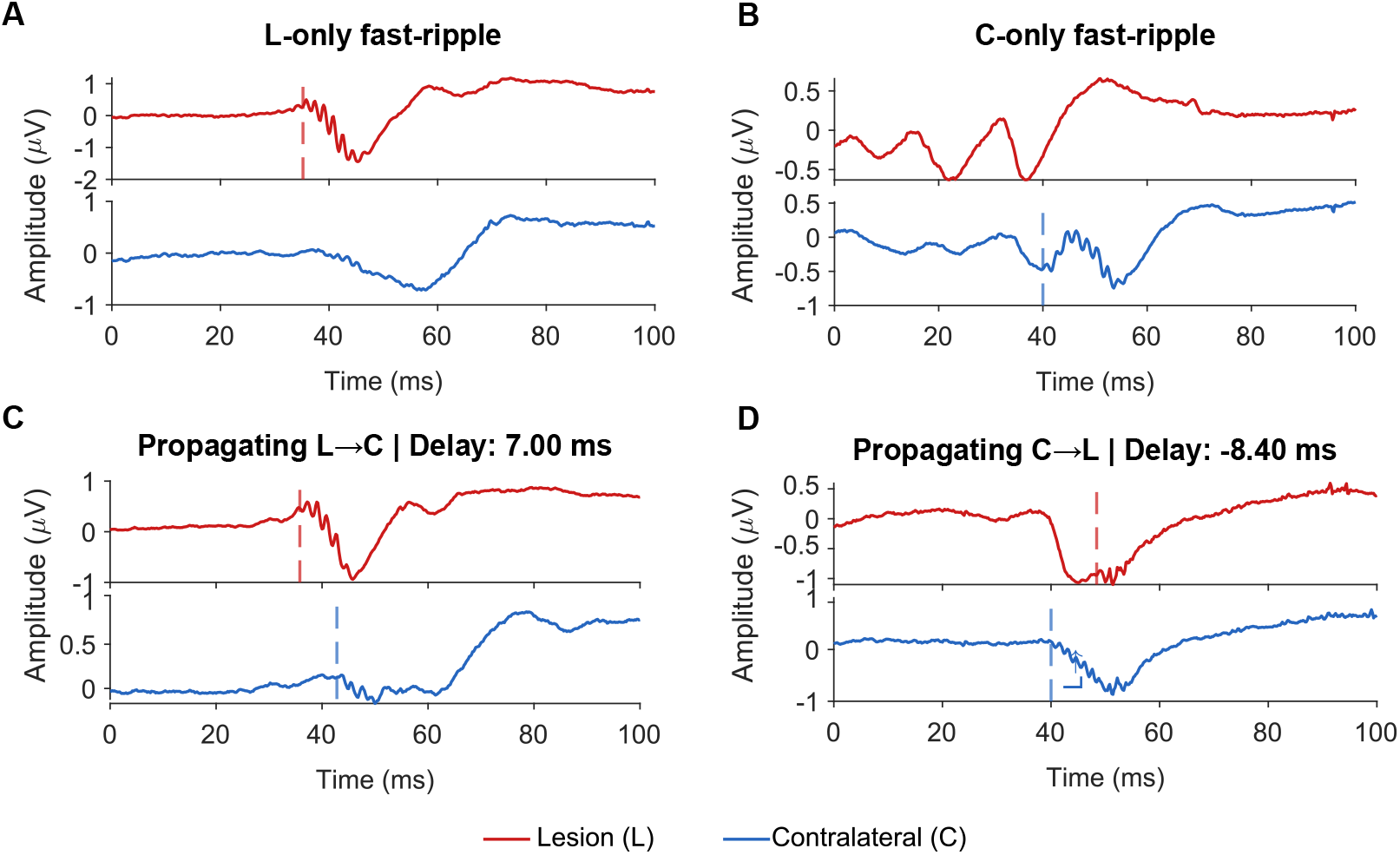
Representative examples of fast-ripple events illustrating the four hemispheric categories. (A) A Lesion-only fast-ripple, visible as a discrete oscillation in the lesional trace with no corresponding event in the contralateral trace. (B) A C-only fast ripple, confined to the contralateral channel with no corresponding oscillation in the lesional trace. (C) A propagating L→C event, in which a fast ripple detected in the lesional channel is accompanied by a time-lagged oscillation of similar frequency in the contralateral channel. (D) A propagating C→L event, showing the reverse pattern, with the fast ripple originating in the contralateral channel and appearing subsequently in the lesional channel.

Fast ripple events propagated in both directions at comparably low absolute rates across all periods (Fig. 7A), with L→C and C→L rates remaining below ≈ 1 event/h and showing no marked evolution across the three periods. Ripple and gamma propagation was observed in only one animal during the early-epileptic and late-epileptic periods (data not shown). We next classified fast ripples and ripples by their hemispheric generation as either lesion-generated (L-generated) or independent (C-generated). The L-generated fast-ripple rate did not change across periods (Fig. 7B, top; Friedman test, *p* = 0.41, *W* = 0.11, *n* = 7), whereas the rate of independently generated contralateral fast ripples increased (Fig. 7B, bottom; Friedman test, *p* = 0.01, Kendall’s *W* = 0.58, *n* = 7), with the pre- to early-epileptic contrast reaching significance on post-hoc comparison (Wilcoxon signed-rank, *p* = 0.008, *r* = 0.63). Ripples showed the similar pattern: the L-generated rate did not change (Fig. 7C, top; Friedman test, *p* = 0.07, Kendall’s *W* = 0.32, *n* = 7), while the independently generated contralateral rate increased from the pre- to the early-epileptic period (Fig. 7C, bottom; Friedman *p* = 0.012, Kendall’s *W* = 0.633, Wilcoxon signed-rank, *p* = 0.039, *r* = 0.53). For fast-ripples the L-only proportion gradually decreased from the pre-epileptic (Fig. 7D, 73.4%, *n* = 8), to the early-epileptic (Fig. 7E, 58.9%, *n* = 8), to the late-epileptic (Fig. 7F, 37.7%, *n* = 7) period (Friedman test, *p* = 0.005, Kendall’s *W* = 0.67, *n* = 7), and the proportion generated in the contralateral hemisphere increased correspondingly (Friedman test, *p* = 0.021, Kendall’s *W* = 0.48). For both measures, with the post-hoc tests, the pre- to late-epileptic period showed significance (Wilcoxon signed-rank, L-only *p* = 0.023, *r* = 0.63, *n* = 7; C-only *p* = 0.047, *r* = 0.60, *n* = 7), whereas the adjacent-period comparisons did not (all *p* 0.078). The proportions of ripples shifted in the same direction (Fig. 7G-I; 74.0%, 64.6% and 51.1%), but this change was not statistically supported (Friedman test, *p* = 0.20, Kendall’s *W* = 0.20; all post-hoc *p* 0.22). Directional L→C propagation remained below 3% of events in all phases for both fast-ripples and ripples, indicating that this redistribution arose from the independent generation of events in the contralateral hemisphere rather than from increased active spread out of the lesion. Consistent with the channel-level recruitment reported in Section 3.1, the epileptiform network thus shifted from a focal, lesion-confined organization before the first spontaneous seizure toward bilaterally independent activity in the chronic period.

**Figure 7:**
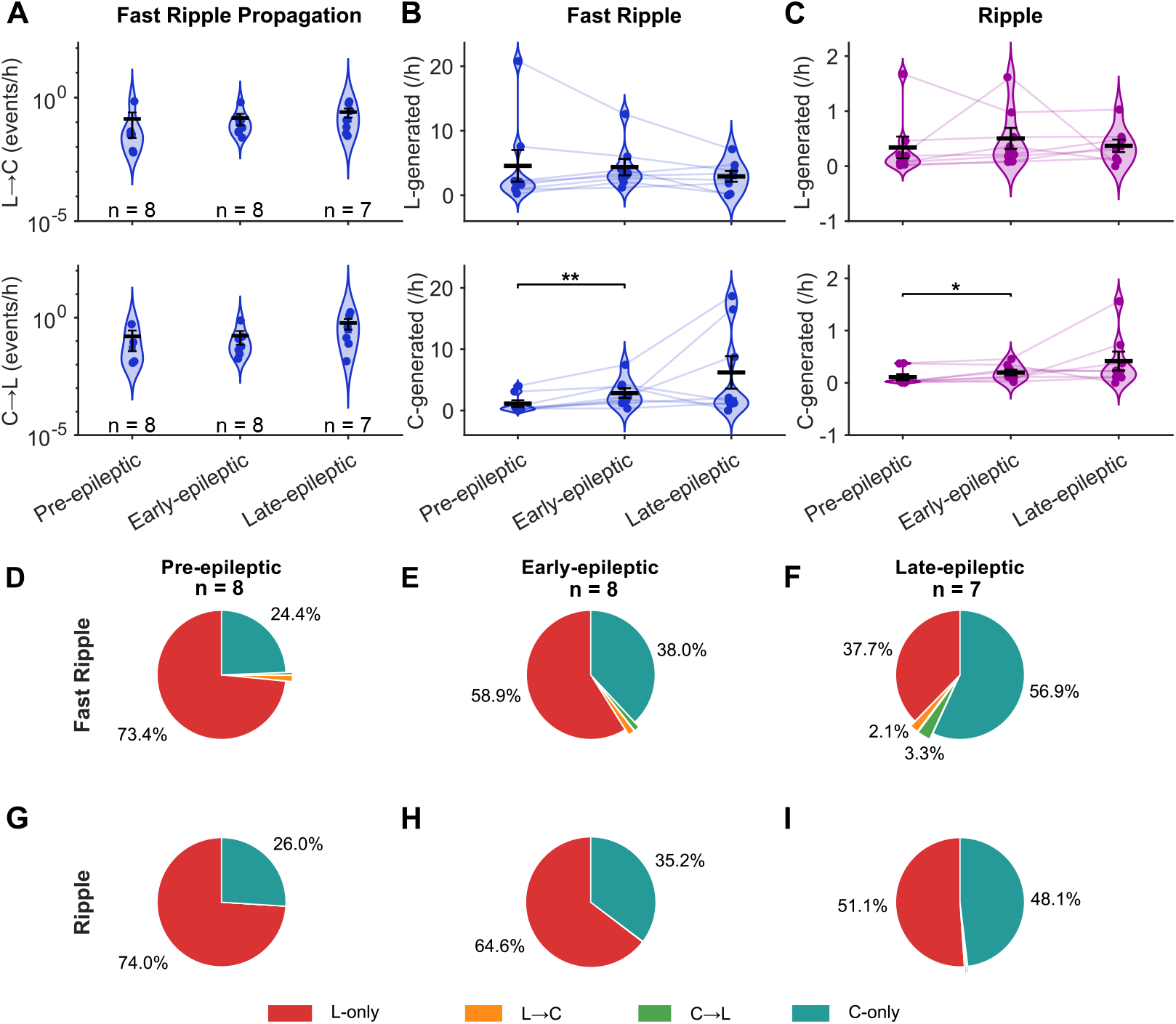
HFO propagation dynamics and spatiotemporal evolution. (A) Propagation rates (events/h) for lesion-to-contralateral (L→C) and contralateral-to-lesion (C→L) directions for fast ripple do not demonstrate facilitated propagation from the lesion or vice versa from the contralateral neocortex. (B, C) Hourly rates of independently generated fast ripples (B) and ripples (C) in the lesional (top; L-generated) and contralateral (bottom; C-generated) channels. The rate of lesion-generated events did not change across periods for either subtype, whereas the rate of independently generated contralateral events increased, reaching significance between the pre- and early-epileptic periods. Violins show the distribution across animals, circles individual animals connected across periods. (D–I) Proportions of fast ripple (D–F) and ripple (G–I) events classified by hemispheric origin during the pre-epileptic (D, G), early-epileptic (E, H) and late-epileptic (F, I) periods. The proportion of events confined to the lesion decreased for both subtypes while the proportion generated in the contralateral hemisphere increased, and events propagating in either direction remained a minor fraction throughout. n is the number of animals.

## 4. Discussion

This study demonstrates that changes in the spatiotemporal properties of HFOs mark the progression of FCD II-related epilepsy and the progressive recruitment of extralesional cortical regions into the epileptic network. Our findings further demonstrate that observations previously derived from TLE models have analogous pathophysiological relevance to neocortical forms of focal epilepsy. The principal finding is that, whereas the lesional cortex maintained chronically elevated but largely stable HFO activity after epilepsy onset, the contralateral cortex exhibited a progressive increase in all HFO subtypes, indicating continuing expansion and reorganization of the epileptic network beyond the primary lesion.

Experimental evidence that HFOs can track epileptogenesis and subsequent epilepsy progression has been derived predominantly from models of TLE. Seminal studies by Bragin and colleagues demonstrated that pathological HFOs emerge early after an epileptogenic insult, before the appearance of spontaneous seizures, and preferentially in animals that subsequently develop epilepsy (Bragin et al., 2000, 2004). These observations led to the concept that pathological HFOs reflect the formation of small pathological neuronal clusters that progressively develop during epileptogenesis and may subsequently recruit interconnected regions into an expanding epileptic network. Importantly, not only HFO occurrence but also their spectral and temporal properties evolve during epileptogenesis, with initially brief, lower-frequency oscillations developing into longer events containing progressively higher frequencies (Jones et al., 2015). Longitudinal studies in the pilocarpine model further demonstrated dynamic changes in both the rate and spatial distribution of HFOs during the transition from the latent to the chronic epileptic state (Salami et al., 2014; Lévesque et al., 2021). For example, HFOs associated with IEDs in the entorhinal cortex increased before the first spontaneous seizure, while IEDs with superimposed fast ripples become particularly prominent during established epilepsy and their occurrence correlated with subsequent seizure activity (Lévesque et al., 2011; Salami et al., 2014; Behr et al., 2015). Moreover, epileptogenesis is accompanied by redistribution of HFO activity between interconnected limbic structures, suggesting that their spatiotemporal evolution reflects progressive reorganization rather than merely increasing excitability within a fixed epileptic focus. Consistent with this concept, remote regions can eventually generate pathological HFOs independently of the primary focus, providing evidence for the development of self-sustained large-scale epileptic networks (Sheybani et al., 2018). Collectively, these studies indicate that HFOs, and particularly fast ripples, are not static markers of epileptogenic tissue but dynamic electrophysiological signatures of the development, reorganization, and expansion of epileptic networks. In our study, the temporal dynamics of the HFO subtypes differed substantially between the two hemispheres. In the lesional cortex, none of the HFO subtypes displayed a day-wise difference from the pre-epileptic phase, while the contralateral cortex displayed an increase in all four event types over the post-epileptic period, with fast ripples changing earliest and remaining elevated for the remainder of the recording period, whereas gamma and ripple rates shifted only later. This pattern indicates that the contralateral hemisphere, rather than the lesion, carries the time-dependent signature of disease progression, and that fast ripples are the first subtype to manifest this contralateral recruitment.

These changes had a clear spatiotemporal structure rather than affecting the limbic network as a whole. Ripples are essentially absent from the dentate gyrus in healthy animals, yet they appear there while epilepsy is still developing, in both kainic acid- and pilocarpine-treated rats (Bragin et al., 2004). After epilepsy is established, the rate of interictal spikes carrying ripples keeps rising in the dentate gyrus, while CA3, the entorhinal cortex and the subiculum show no such increase (Lévesque et al., 2011). Salami et al. (2014) reported a comparable regional split for one class of spike waveform: events with ripples or fast ripples became less frequent in the entorhinal cortex and more frequent in CA3 once the chronic phase began. A recent bilateral recording study in the intrahippocampal kainic acid model points in the same direction, with interictal spikes rising most steeply in the dentate gyrus on the injected side and fast ripples increasing more than ripples in that hemisphere (Friscourt et al., 2026). While TLE models have shown that HFOs evolve dynamically during epileptogenesis in the primary epileptogenic focus, our data show that in FCDII-related epilepsy, this dynamic modulation occurs only in the contralateral cortex. This divergence is biologically plausible. In post-status epilepticus TLE models, epileptogenesis proceeds from an initially intact hippocampus through progressive cell loss and network reorganization (Bragin et al., 1999; Lévesque et al., 2011). The pre-epileptic HFO rise may therefore reflect a dynamic reorganization process with an identifiable onset that can be recorded immediately or just a few days after status epilepticus. In FCDII, by contrast, the dysplastic lesion is established during embryonic brain development, long before recording begins, and the cortex within and surrounding it is structurally and functionally abnormal from the outset (Chvojka et al., 2024) and, thus, may display a saturated pathological baseline with a chronically elevated HFO activity (Bragin et al., 2010). The lesional channel may thus already be operating near its ceiling of pathological activity at the start of monitoring, leaving little dynamic range for a detectable seizure-related escalation.

The propagation analysis allowed us to distinguish between two mechanistically distinct explanations for the rising contralateral HFO burden: increased spread of activity out of the lesion, or independent HFO generation within the contralateral hemisphere. The data strongly favour the latter. Directional L→C propagation accounted for less than 3% of fast ripple and ripple events in all three periods and showed no temporal evolution, ruling out a propagation-driven explanation. Instead, the fraction of events arising from the contralateral hemisphere (C-only and C→L) increased across analyzed disease phases, reflecting genuine *de novo* generation of HFOs in the contralateral cortex. This pattern raises the question of what mechanism drives the progressive establishment of an autonomous contralateral HFO generator.

One possibility is secondary epileptogenesis, in which repeated seizure activity from the lesional hemisphere induces lasting changes in contralateral cortical excitability (Jiruska et al., 2023). In the present model, 95.7% of seizures originated from the lesional hemisphere, providing a sustained and lateralized drive that could progressively modify synaptic strength and network connectivity in homotopic contralateral regions. If pathological HFOs indeed mark tissue capable of generating seizures, the progressive emergence of independently generated HFOs in the contralateral cortex raises the possibility that, at later stages of disease, these regions may acquire sufficient epileptogenicity to become additional or even alternative seizure onset zones independent of the FCD lesion (Proietti Onori et al., 2021).

It is well established that chronic epilepsy can be associated with the progressive spread of epileptic activity beyond the primary epileptic focus, resulting in the gradual conversion of connected brain regions into epileptogenic tissue through the mechanisms related to kindling (Morimoto et al., 2004; Goddard, 1967). Both recurrent seizures and persistent interictal activity have been implicated in this process. Based on experimental and clinical observations in drug-resistant focal epilepsy ongoing epileptic activity induces epileptogenic reorganization in anatomically connected regions, leading to the formation of secondary, and eventually tertiary, epileptic foci that can generate epileptic activity independently of the primary focus (Shen et al., 2021; Morrell, 1959b,a). The formation of secondary epileptic foci may result from the combined effects of recurrent seizures and continuous exposure of projection areas to interictal epileptiform activity. In particular, seizures and interictal discharges containing high-frequency oscillations (HFOs), which reflect highly synchronized action potential firing, are expected to exert a particularly strong influence on downstream networks. Repeated barrages of excessive synchronous activity can progressively modify distant circuits through mechanisms resembling kindling and activity-dependent synaptic remodeling (Bragin et al., 2000; Jiruska et al., 2017).

These processes are thought to involve sustained NMDA receptor activation, calcium-dependent intracellular signaling, altered gene expression, and long-term structural and functional synaptic changes (Esclapez et al., 1999; Stasheff et al., 1989). Compelling experimental evidence for seizure-induced remodeling of projection areas was provided by Khalilov et al. (2003). Using an in vitro bilateral hippocampal preparation, they demonstrated that repeated seizures originating in one hippocampus consistently propagated to the contralateral side and eventually generated a secondary epileptic focus capable of producing spontaneous seizures independently of the primary hippocampus, even after pharmacological or anatomical blockade of commissural fibers. Importantly, this epileptogenic transformation required intact glutamatergic transmission, as NMDA receptor antagonists prevented the development of the secondary focus. Similar network remodeling has been demonstrated following repeated focal activation with glutamate or NMDA, providing further evidence that excessive excitatory activity alone is sufficient to induce long-lasting epileptogenic changes (Croucher et al., 1995; Croucher and Bradford, 1989). Likewise, intracellular recordings from neocortical projection areas have demonstrated a functional transformation of principal neurons, which shift from regular spiking to burst-firing behavior resembling neurons within the primary epileptic focus (Brener et al., 1991). This concept is highly relevant to our FCD model, in which projection areas are known to exhibit both morphological and functional abnormalities before the appearance of overt epileptic activity. Experimental models of FCD induced by mTOR or RHEB mutation have shown that callosal projections from the lesion fail to terminate selectively within cortical layers II/III and instead are distributed throughout the cortical column (Proietti Onori et al., 2021; Procházková et al., 2025). Moreover, excitatory presynaptic terminals are enlarged in both the lesion and the contralateral cortex, including giant synapses with diameters up to five times larger than normal (Proietti Onori et al., 2021; Procházková et al., 2025). Intracellular recordings further demonstrated that principal neurons in the contralateral cortex exhibit abnormal burst firing and enhanced excitatory synaptic transmission, both of which promote network hyperexcitability. These observations provide a mechanistic explanation for the contralateral cortex being so permissive for the development of independent epileptiform activity. We propose that this transition results from the combined effects of activity-dependent epileptogenic remodeling superimposed on pre-existing structural abnormalities of the contralateral cortex in FCD.

## 5. Conclusion

Longitudinal spatiotemporal analysis of IEDs and pathological HFOs provides unique insight into the dynamic reorganization of FCD-related epileptic networks during disease progression. Although the dysplastic lesion remained the principal seizure onset zone throughout the study, the temporal evolution of these biomarkers revealed progressive functional recruitment of the contralateral cortex. Importantly, this reorganization was characterized by the emergence of independent HFO generators rather than increased interhemispheric propagation, demonstrating that longitudinal biomarker analysis can distinguish between network propagation and network remodeling. Our findings also have important clinical implications. Focal cortical dysplasia should not be viewed as an isolated structural lesion but rather as the center of a progressively reorganizing epileptic network. This concept is particularly relevant for presurgical evaluation, as the functional extent of the epileptic network may extend beyond the radiologically or histologically defined lesion because of pre-existing connectivity abnormalities and progressive network remodeling. Whether independently generated epileptiform activity in distant cortical regions ultimately contributes to postsurgical seizure recurrence or surgical failure remains to be determined in future experimental and clinical studies.

## 6. Funding

This study was supported by grants of the Czech Science Foundation [25-17580S], the Ministry of Health of the Czech Republic [NW24-08-00394, NW26-04-00517], the Ministry of Education Youth and Sports of the Czech Republic [EU – Next Generation EU: LX22NPO5107], ERDF-Project Brain dynamics [CZ.02.01.01/00/22_008/0004643], and the Charles University project EXCITE [UNCE24/MED/021] and PRIMUS [23/MED/011]. Access to CESNET storage facilities provided by the project „e-INFRA CZ“ under the program „Projects of Large Research, Development, and Innovations Infrastructures” [LM2018140].

## 7. Author contributions

P.J. and J.K. contributed to the conception and design of the study; N.K., M.S., N.P., B.H., M.K., O.N., H.P., J.K., and P.J. contributed to experimental work or data analysis. N.K., J.K., and P.J. contributed to drafting the text or preparing the figures.

## 8. CRediT authorship contribution statement

Nedime Karakullukcu: Formal analysis, Visualization, Validation, Methodology, Writing – original draft, Writing – review & editing. Michal Scheibel: Resources, Data curation. Natalie Prochazkova, Bohdana Hruskova, Michaela Kralikova, Ondrej Novak, and Helena Pivonkova: Data curation, Methodology. Jan Kudlacek: Data curation, Formal analysis, Investigation, Methodology, Supervision, Writing – original draft, Writing – review & editing. Premysl Jiruska: Conceptualization, Formal analysis, Funding acquisition, Methodology, Project administration, Supervision, Writing – original draft, Writing – review & editing.

## 9. Declaration of competing interest

No conflicts of interest to disclose.

## 10. Data availability

Recorded data and analytical tools used in this study are available from the corresponding author upon request.

## 11. Appendix A. Supplementary material

## Notes

### Competing Interest Statement

The authors have declared no competing interest.

