## Supplementary Material for "High-frequency oscillations reveal progressive recruitment of remote cortex into the epileptic network in a mouse model of focal cortical dysplasia type II"

### 1. Supplementary Methods

#### *1.1. Feature Extraction and Data Preparation*

Training data were obtained from four animals, including two animals from the current dataset and two independent animals that were not included in the longitudinal analyses because their seizure profiles did not provide sufficient pre-epileptic period data before the first spontaneous seizure. These exclusions were based solely on data availability and were not related to high-frequency oscillations (HFO) characteristics. In total, 23,890 putative events were manually labeled as HFO or non-HFO. Within each animal the number of non-HFO events was matched to the number of HFO events, giving 11,945 events per class. A total of 17 distinct features were extracted from each candidate HFO event to characterize its morphology and spectral properties. The features are described in Supplementary Table 1.

#### *1.2. Classification and Validation Strategy*

The feature set was used to train a binary classification model. The classifier was trained on data from four animals (N = 4) using leave-one-subject-out cross-validation (LOSOCV) to ensure subject-independent generalisation. In each fold, the model was trained on three subjects and tested on the remaining unseen subject, preventing subject-specific data leakage (Saeb et al., 2017). An ensemble bagging architecture ($fitcensemble$, method $Bag$, 100 learning cycles, decision-tree weak learners, $MaxNumSplits = 10$) was chosen to enhance robustness against the variability of HFO morphology across different subjects (Breiman, 1996; Mohammed and Kora, 2023). The trained model classified each candidate event as HFO or non-HFO (Supplementary Fig. 1 and Supplementary Fig. 2).

The overall diagnostic ability of the ensemble bagging classifier was further evaluated by receiver operating characteristic (ROC) curve analysis. The ROC curve plots sensitivity (true positive rate (TPR)) against $1 - specificity$ (false positive rate (FPR)) across the full range of decision thresholds.

The area under the ROC curve (AUC) was 0.83 under LOSOCV, indicating that discriminative ability was retained when the classifier was applied to animals that had not contributed to training. The operating threshold was chosen by maximising the difference between the true positive rate and the false positive rate (Hassanzad and Hajian-Tilaki, 2024). This point is marked in red in Supplementary Fig. 3. At this operating point, TPR was 0.88 at a FPR of 0.31, corresponding to a specificity of 0.69.

The spread of the fold-wise ROC curves ($0.81 \pm 0.06$ across the four folds) indicates that classification performance did not depend strongly on which animal was held out.

Representative raw traces illustrating the four categories of fast ripple events defined in the Methods of the main manuscript are shown in Supplementary Fig. 4.

Supplementary Table 1: Summary of extracted features, descriptions, formulas, and explanations of symbols. A total of 17 features were used.

| **Feature** | **Description** | **Formula** | **Explanation** |
|---|---|---|---|
| Kurtosis | Measures the peakedness of the amplitude distribution. | $\text{kurt}(x) = \frac{1}{N}\sum\left(\frac{x-\mu}{\sigma}\right)^4$ | $x$ = signal amplitude, $\mu$ = mean, $\sigma$ = standard deviation, $N$ = number of samples |
| Skewness | Quantifies the asymmetry of the distribution. | $\text{skew}(x) = \frac{1}{N}\sum\left(\frac{x-\mu}{\sigma}\right)^3$ | Same as above |
| Shannon Entropy | Measures signal complexity. | $H = -\sum p_i \log(p_i)$ | $p_i$ = probability of the i-th amplitude bin |
| Log Energy Entropy | Reflects energy distribution. | $H_{\text{logE}} = \sum \log(x^2)$ | $x$ = signal amplitude |
| Energy | Total signal energy in the event window. | $E = \sum x^2$ | $x$ = signal amplitude |
| DWT Coefficients | Time–frequency localized features via wavelets. | $W_{j,k} = \langle x(t), \psi_{j,k}(t)\rangle$ | $x(t)$ = signal, $\psi_{j,k}$ = wavelet at scale $j$ and shift $k$ |
| Delta Power (0.5–4 Hz) | Power in delta band. | $P_\delta = \int_{0.5}^{4} \lvert X(f)\rvert^2 df$ | $X(f)$ = Fourier transform of signal |
| Theta Power (4–8 Hz) | Power in theta band. | $P_\theta = \int_{4}^{8} \lvert X(f)\rvert^2 df$ | $X(f)$ = Fourier transform of signal |
| Delta/Theta Ratio | Normalizes low-frequency changes. | $\frac{P_\delta}{P_\theta}$ | $P_\delta$ = delta power, $P_\theta$ = theta power |
| Alpha Power (8–12 Hz) | Power in alpha band. | $P_\alpha = \int_{8}^{12} \lvert X(f)\rvert^2 df$ | $X(f)$ = Fourier transform of signal |
| Beta Power (13–30 Hz) | Power in beta band. | $P_\beta = \int_{13}^{30} \lvert X(f)\rvert^2 df$ | $X(f)$ = Fourier transform of signal |
| Alpha/Beta Ratio | Relationship between alpha and beta activity. | $\frac{P_\alpha}{P_\beta}$ | $P_\alpha$ = alpha power, $P_\beta$ = beta power |
| Gamma+Ripple Power (30–250 Hz) | Combined high-frequency content. | $P_{\gamma+r} = \int_{30}^{250} \lvert X(f)\rvert^2 df$ | $X(f)$ = Fourier transform of signal |
| Fast Ripple Power (250–800 Hz) | Ultrafast high-frequency content. | $P_{\text{FR}} = \int_{250}^{800} \lvert X(f)\rvert^2 df$ | $X(f)$ = Fourier transform of signal |
| Gamma+Ripple / Fast-Ripple Ratio | Relative distribution of high-frequency energy. | $\frac{P_{\gamma+r}}{P_{\text{FR}}}$ | $P_{\gamma+r}$ = gamma+ripple power, $P_{\text{FR}}$ = fast ripple power |
| All-Band Power (0.5–1000 Hz) | Full-band energy measure. | $P_{\text{all}} = \int_{0.5}^{1000} \lvert X(f)\rvert^2 df$ | $X(f)$ = Fourier transform of signal |
| AR Coefficients | Captures temporal structure via autoregressive modeling. | $x(t) = \sum_{i=1}^{p} a_i x(t-i) + e(t)$ | $x(t)$ = signal at time $t$, $a_i$ = AR coefficients, $p$ = model order, $e(t)$ = error term |

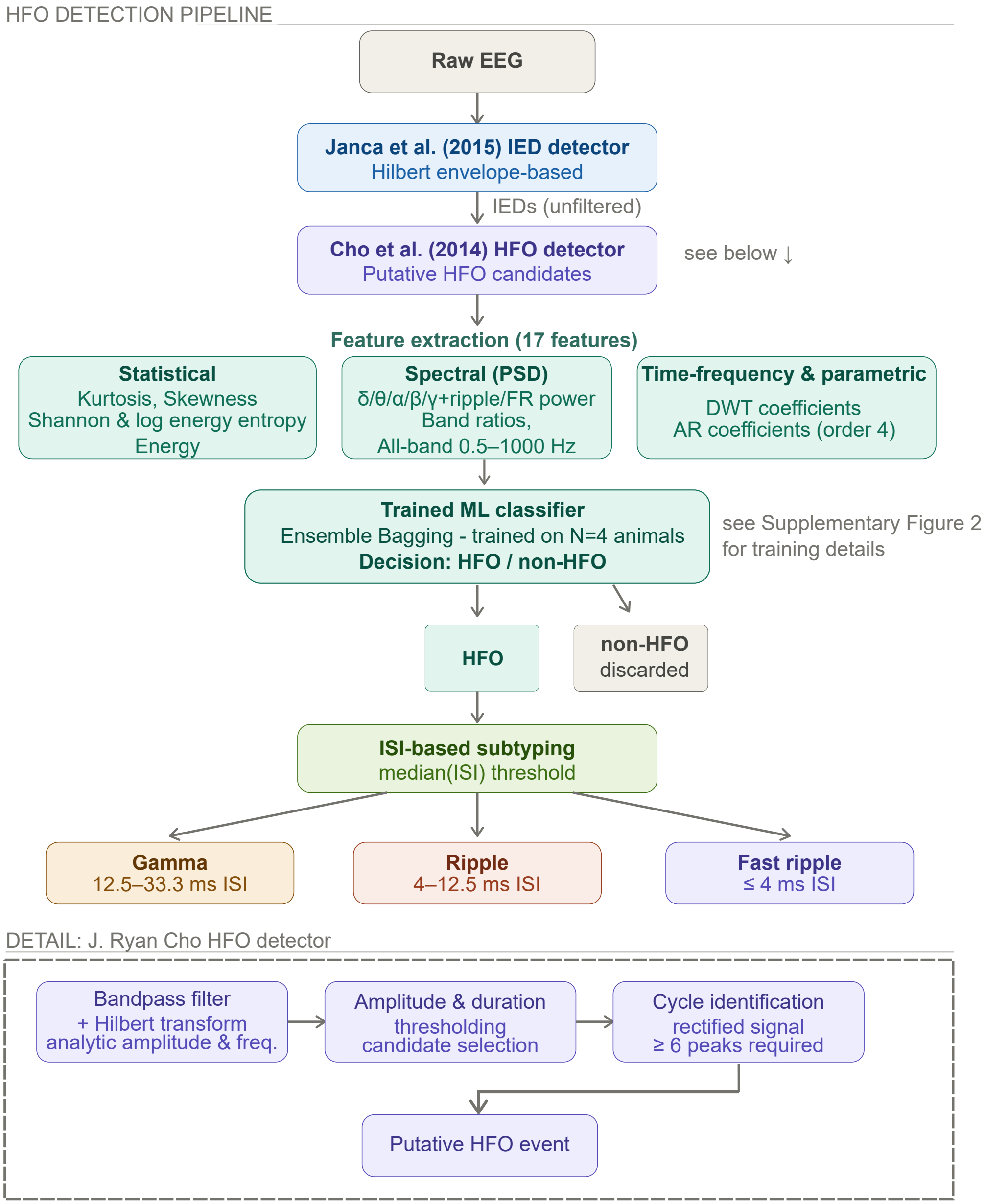


Supplementary Figure 1: Overview of the HFO detection pipeline, from raw EEG through interictal epileptiform discharges (IED) detection, candidate HFO detection, feature extraction and classification, to subtyping of accepted events into gamma, ripple and fast ripple. The dashed box details the candidate detection step. Full descriptions are given in the Methods of the main manuscript. ISI, inter-spike intervals (Jiruska et al., 2010); PSD, power spectral density; DWT, discrete wavelet transform; FR, fast ripple; ML, machine learning; AR, autoregressive.

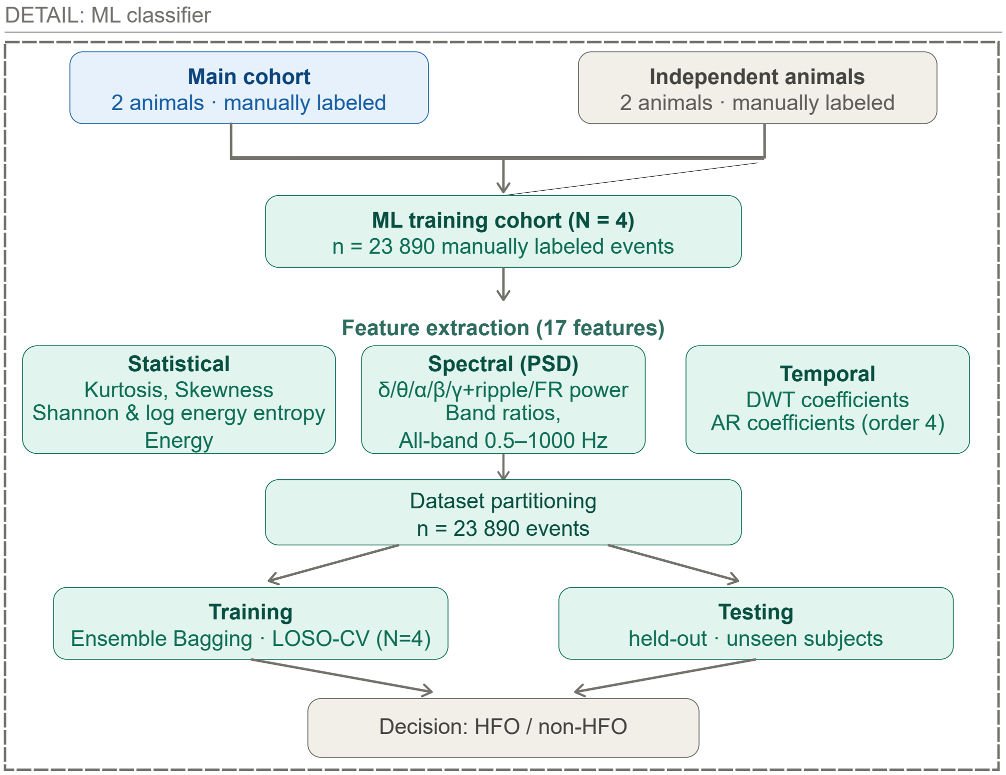


Supplementary Figure 2: Training and validation of the machine learning classifier. Manually labeled events from four animals, two from the cohort analyzed here and two independent animals, formed a training set of 23,890 events. An ensemble bagging classifier was trained using leave-one-subject-out cross-validation across the four animals.

### 2. Supplementary Results

#### *2.1. Machine Learning Performance*

The LOSOCV results demonstrated that the ensemble bagging-based classifier considerably improved the HFO detection performance compared to the conventional Hilbert-based HFO detector adapted from Cho et al. (2014), which is a widely used rule-based baseline. When applied to the same labeled dataset, the conventional detector achieved an accuracy of 0.71, precision of 0.65, TPR of 0.83, and an $F1$ score of 0.73, whereas the proposed machine learning (ML)-based pipeline yielded higher accuracy (0.78), precision (0.74), TPR (0.88), and $F1$ score (0.80), together with an AUC of 0.83. The proposed pipeline achieved both a higher true positive rate and a lower false positive rate than the rule-based detector, indicating that the improvement reflects better discrimination rather than a shift of the decision threshold. While both approaches showed comparable sensitivity, the improved balance between sensitivity and specificity achieved by the ML-based pipeline supports its use for reproducible and scalable quantification of HFO dynamics across extended recordings.

The ReliefF algorithm was employed to rank the contribution of each feature to the classification task (Robnik-Šikonja and Kononenko, 2003). The four most discriminative features were the gamma+ripple / fast-ripple ratio, gamma+ripple power (30–250 Hz), beta power (13–30 Hz), and alpha power (8–12 Hz). The gamma+ripple/fast-ripple ratio and the gamma+ripple power are directly related to the core characteristics of HFOs. The

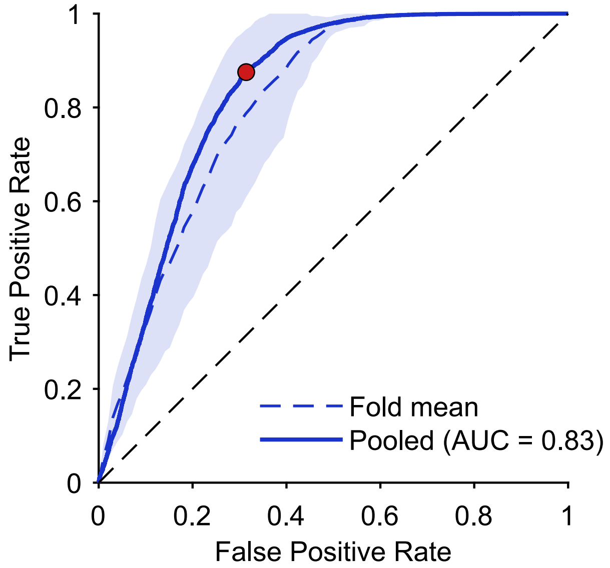


Supplementary Figure 3: Receiver operating characteristic (ROC) curve for leave-one-subject-out (LOSOCV) cross-validated HFO classification. The ensemble classifier (bagged decision trees) was trained on data from three subjects and tested on the held-out fourth, repeated for all four subjects (folds). The solid line shows the pooled ROC curve, computed by concatenating the held-out predictions from all folds into a single dataset (AUC = 0.832). The dashed line shows the fold-mean ROC curve, obtained by computing each fold's own ROC curve separately, interpolating it onto a common false-positive-rate grid, and averaging across the four folds; the shaded band shows $\pm 1$ SD across folds, reflecting subject-to-subject variability in classifier performance. The red marker indicates the operating point maximizing Youden's J statistic (sensitivity - [1 - specificity]) on the pooled curve.

prominence of the gamma+ripple power (30–250 Hz) is consistent with ripple-band energy being the dominant discriminative signal. The ratio between these bands and fast ripples may also help to separate genuine oscillations from broadband artefacts. The inclusion of beta (13–30 Hz) and alpha (8–12 Hz) power bands indicates that background activity in the alpha and beta bands also contributed to the classification.

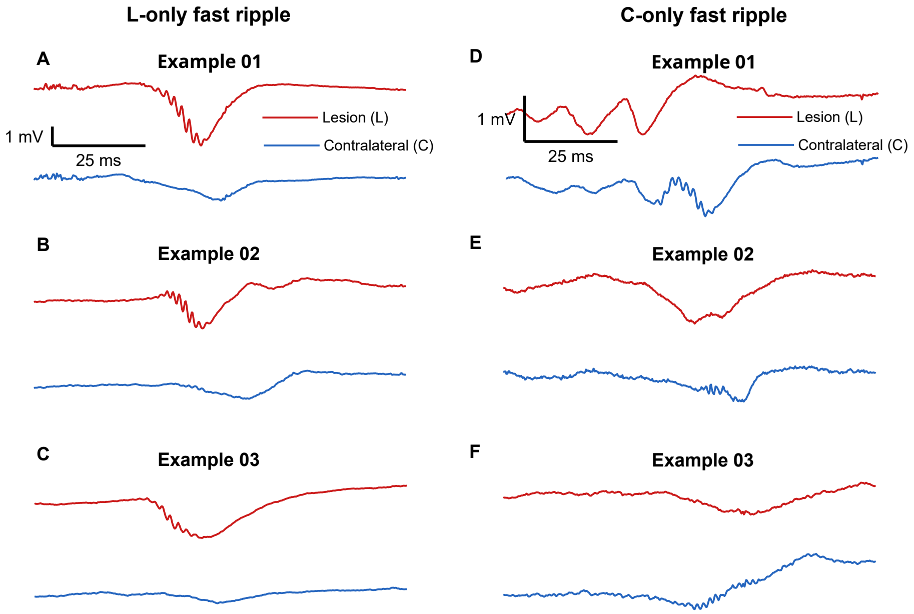


(a) Isolated raw fast ripple events. The left column (A–C) shows three examples of fast ripples detected in the lesional channel only (L-only), with no corresponding event in the contralateral channel, and the right column (D–F) three examples detected in the contralateral channel only (C-only). Scale bars in the top row apply to all panels.

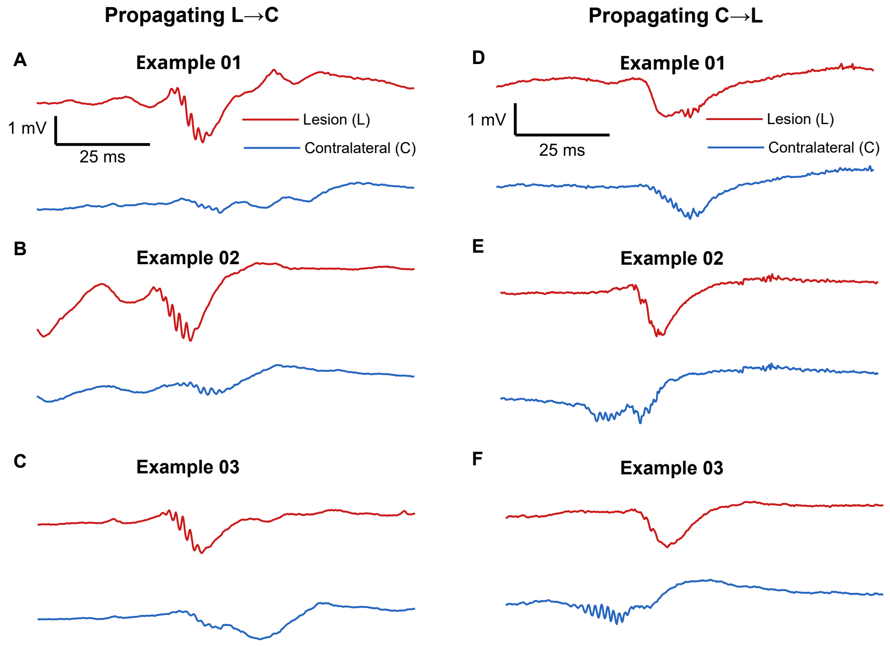


(b) Propagating events, in which the same fast ripple was detected in both channels within the propagation window. The left column (A–C) shows three examples propagating from the lesion to the contralateral cortex (L→C) and the right column (D–F) shows three examples propagating in the opposite direction (C→L). Scale bars in the top row apply to all panels.

Supplementary Figure 4: Representative raw traces of fast ripple events classified by hemispheric generation and propagation. In all panels the red trace was recorded from the lesional channel (L) and the blue trace from the contralateral homotopic channel (C). Examples are single events rather than averages and were taken from different disease periods: L-only events from the pre- and early-epileptic periods, L→C events from the early-epileptic period, and C-only and C→L events from the late-epileptic period. Event classification followed the criteria described in the Methods of the main manuscript.
